# TLR3-TRIF pathway activation by *Neospora caninum* RNA enhances infection control

**DOI:** 10.1101/327999

**Authors:** Vanessa dos Santos Miranda, Flávia Batista Ferreira França, Mylla Spirandelli da Costa, Vanessa Resende Souza Silva, Caroline Martins Mota, Patrício da Silva Cardoso Barros, Kleber Simônio Parreira, Fernanda Maria Santiago, Jose Roberto Mineo, Tiago Wilson Patriarca Mineo

## Abstract

*Neospora caninum* is a protozoan parasite closely related to *Toxoplasma gondii* and has been studied for causing neuromuscular disease in dogs and abortions in cattle. It is recognized as the major cause of economic losses in bovine products. In that sense, this study aimed to evaluate the role of TLR3-TRIF dependent resistance against *N. caninum* infection. We observed that TLR3^−/−^ and TRIF^−/−^ mice presented higher parasite burden, increased inflammatory lesions and reduced production of IL-12p40, TNF, IFN-γ, and NO. Differently from *T. gondii, N. caninum* tachyzoites and its RNA recruited TLR3 and IRF3 to the parasitophorous vacuole (PV). We observed that N. caninum upregulated the expression of TRIF in macrophages, which by its turn upregulated IFN-α and IFN-β in the presence of the parasite. Furthermore, TRIF^−/−^ infected macrophages produced lower levels of IL-12p40 and IFN-α replacement was able to completely restore the production of this key cytokine. Our results have shown that TLR3-TRIF signaling pathway enhances resistance against *N. caninum* infection, since it improves Th1 immune responses that control parasitism and tissue inflammation, which are hallmarks of the disease.

## 1 Introduction

*Neospora caninum* is an obligate intracellular parasite that belongs to the phylum Apicomplexa and is the causative agent of neosporosis. Canids have been described as its definitive hosts, while cattle, sheep, and other warm-blooded species act as its intermediate hosts (1, 2), which can be infected by oral or transplacental routes. *N. caninum* is closely related to *Toxoplasma gondii*, and has been studied in the last decades as a major cause of neuromuscular disease in dogs and repeated abortions in cattle, being recognized as the major identifiable cause of economic losses in beef and dairy industries (3).

The innate immune response triggered by *N. caninum* play an important role in protection of the host, leading to the activation of adaptive responses that restricts parasite proliferation. Innate cells express pattern recognition receptors (PRRs) as the Toll-like receptors (TLRs), that recognize pathogen associated molecular patterns (PAMPs) (4–7). This recognition requires intracellular signaling through adapter molecules, mainly the myeloid differentiation factor 88 (MyD88) or molecule inducing interferon-β with TIR domain (TRIF) (8). Most TLRs share MyD88 as its adapter protein, with exception of TLR3, which only transcribes recognition signals through TRIF, and TLR4, that signals through both pathways depending on the stimulus (9). The MyD88-dependent signaling pathway leads to the activation of MAP kinases (MAPK) and transcription factors as NF-κB, which encode the expression of classic pro-inflammatory mediators. On the other hand, the TRIF-dependent pathway activates Interferon Regulatory Factors (IRFs), resulting in the production of Type I interferon (IFN-β and IFN-α), which may potentiate the production of pro-inflammatory factors such as TNF (8, 10, 11).

Related to *N. caninum* infection, the MyD88-dependent pathways is known to be activated during the infection, acting as a relevant host resistance factor (12, 13). In this context, TLR2 and TLR11 participate in parasite recognition, leading to the activation of antigen presenting cells and polarization of immune responses to a Th1 profile (14, 15). The participation of MyD88-dependent endosomal receptors, such as TLR 7, 8 and 9, has also been suggested during the infection in different animal species (16–18). However, the role of TRIF signaling in the context of *N. caninum* infection is still unclear. It is known that, differently from *T. gondii*, TLR3 recognizes *N. caninum* RNA and induces type I IFN responses (19).

In that sense, this study aimed to evaluate the role of the TLR3-TRIF dependent signaling pathway in resistance to *N. caninum*, using *in vivo* and *in vitro* approaches to observe phenotypes and to elucidate the molecular mechanisms involved in this interaction.

## 2 Results

### 2.1 TRIF is required for proper resistance during the infection by *N. caninum*

In order to verify whether TLR3-TRIF signaling pathway was required for host survival against a severe infection model, we monitored WT, TLR3^−/−^, TRIF^−/−^ and MyD88^−/−^ mice for 30 days after inoculation of *N. caninum* tachyzoites (1×10^7^ parasites/mice). MyD88^−/−^ mice were used as a controls since it has been shown that MyD88-induced signaling is crucial for host resistance to neosporosis (12). WT mice survived the infection, whereas all MyD88^−/−^ mice succumbed after 18 days of inoculation. Regarding the TLR3^−/−^ and TRIF^−/−^ mice, we found a survival rate of 50% and 33%, respectively, demonstrating that this pathway participates in host resistance to infection (Figure 1).

**Figure 1:**
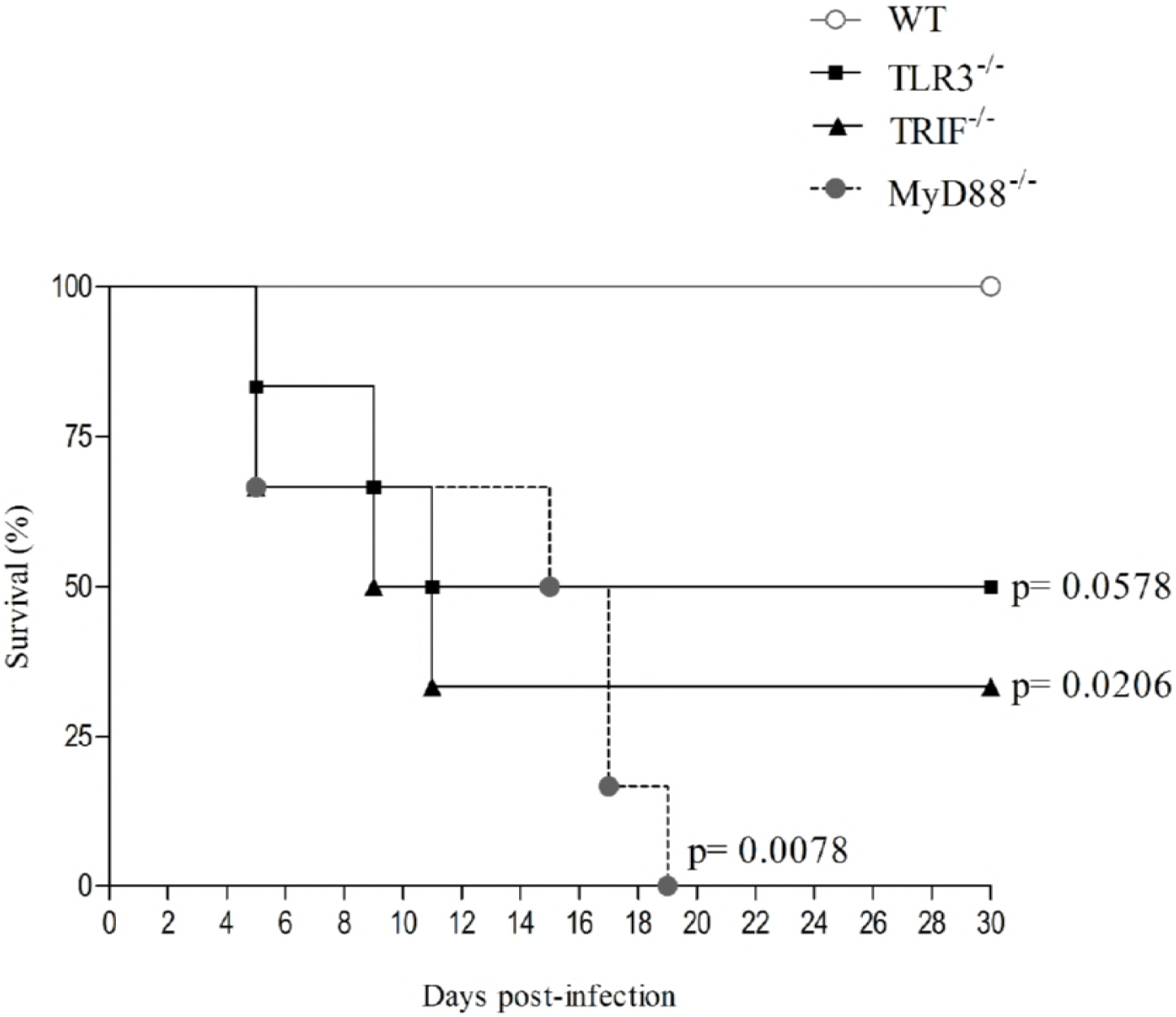
Survival curve of WT, MyD88^−/−^, TLR3^−/−^ and TRIF^−/−^ (n = 5) mice monitored for up to 30 days after infection with 1×10^7^ *N. caninum* tachyzoites. Differences between groups were compared with Kaplan Meier survival analysis, using the log-rank test, with *p* value representing the comparison to WT group. Values are representative of three independent experiments.

Then, in order to evaluate the specific participation of TRIF in control of parasite growth *in vivo*, we infected WT and TRIF^−/−^ mice with a sublethal dose of *N. caninum* tachyzoites (5×10^6^ parasites/mice), for the quantification of parasite genomic DNA copies at distinct phases of infection (Figure 2). We observed that TRIF^−/−^ mice presented increased parasite burden during hyper acute (24 hours of infection, peritoneal cells), acute (7 days of infection; peritoneal cells and lungs), and in the chronic phase (30 days of infection; brain), if compared to the infected WT group. No significant differences in parasite load were observed in the liver (acute phase) between the analyzed groups.

**Figure 2:**
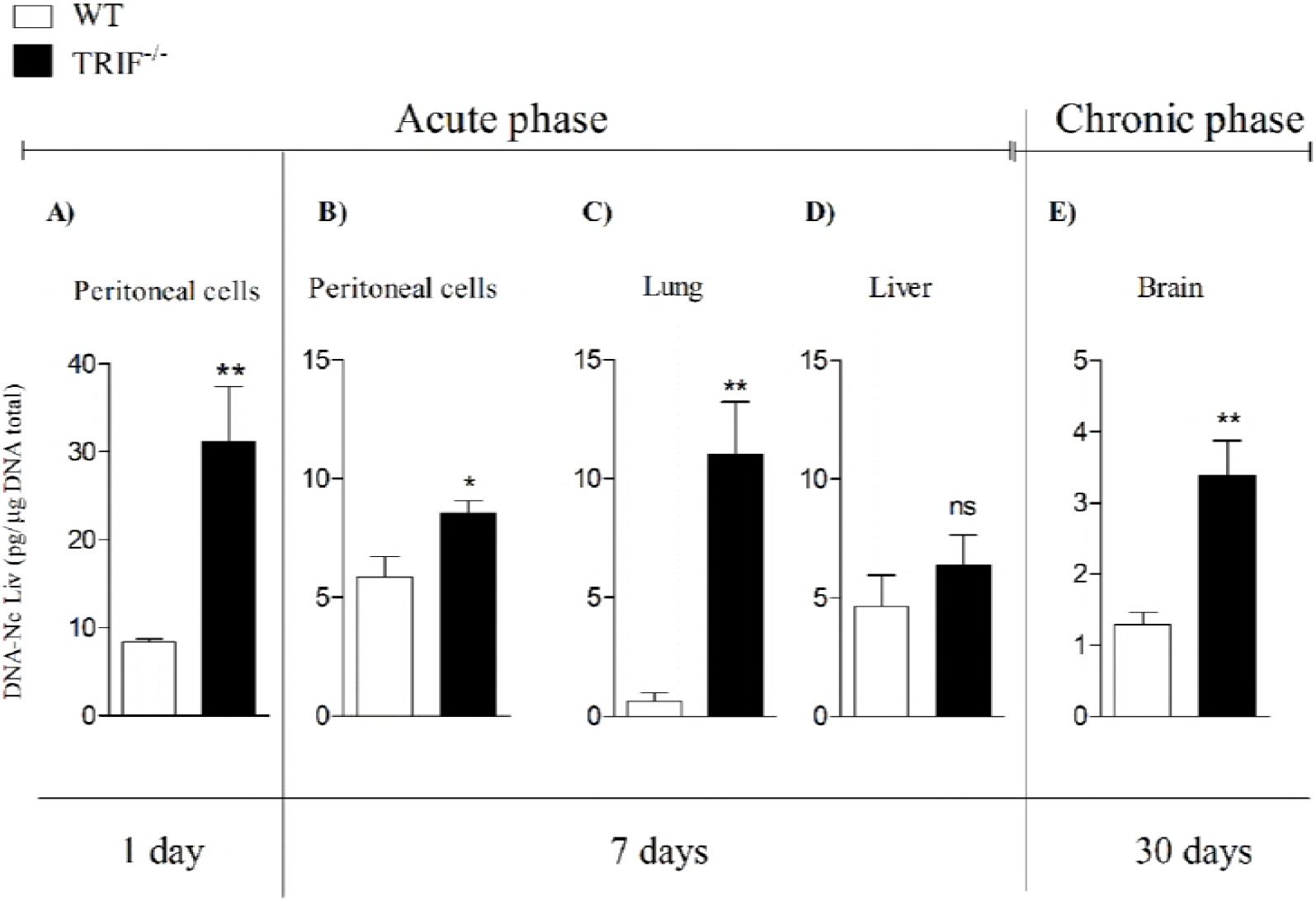
TRIF restricts parasite growth *in vivo*. WT and TRIF^−/−^ mice were infected with 5×10^6^ tachyzoites of *N. caninum* and sacrificed for collection of peritoneal cells, lung, liver and brain tissue in the acute and chronic phases of the infection. The data were obtained from qPCR by amplification of Nc5 gene. Values are expressed as mean ± standard error of the mean (SEM). Data were analyzed using the Mann Whitney test. (* P <0.1 ** P <0.01, ns: not statistically significant). Values are representative of two independent experiments.

Based on these findings, we evaluated the role of TRIF in controlling tissue injury and inflammatory lesions in the different tissues of the infected mice. Histological sections revealed pronounced lesions in the lungs of the infected mice at 7 days post infection, notably worsened in the absence of the TRIF, which presented a significant loss of tissue structure induced by extensive regions of diffuse inflammatory infiltrates. Inflammatory foci were also observed in liver and brain samples from WT and TRIF^−/−^ mice, however no differences in the inflammatory profile were noted between the analyzed groups (Figure 3).

**Figure 3:**
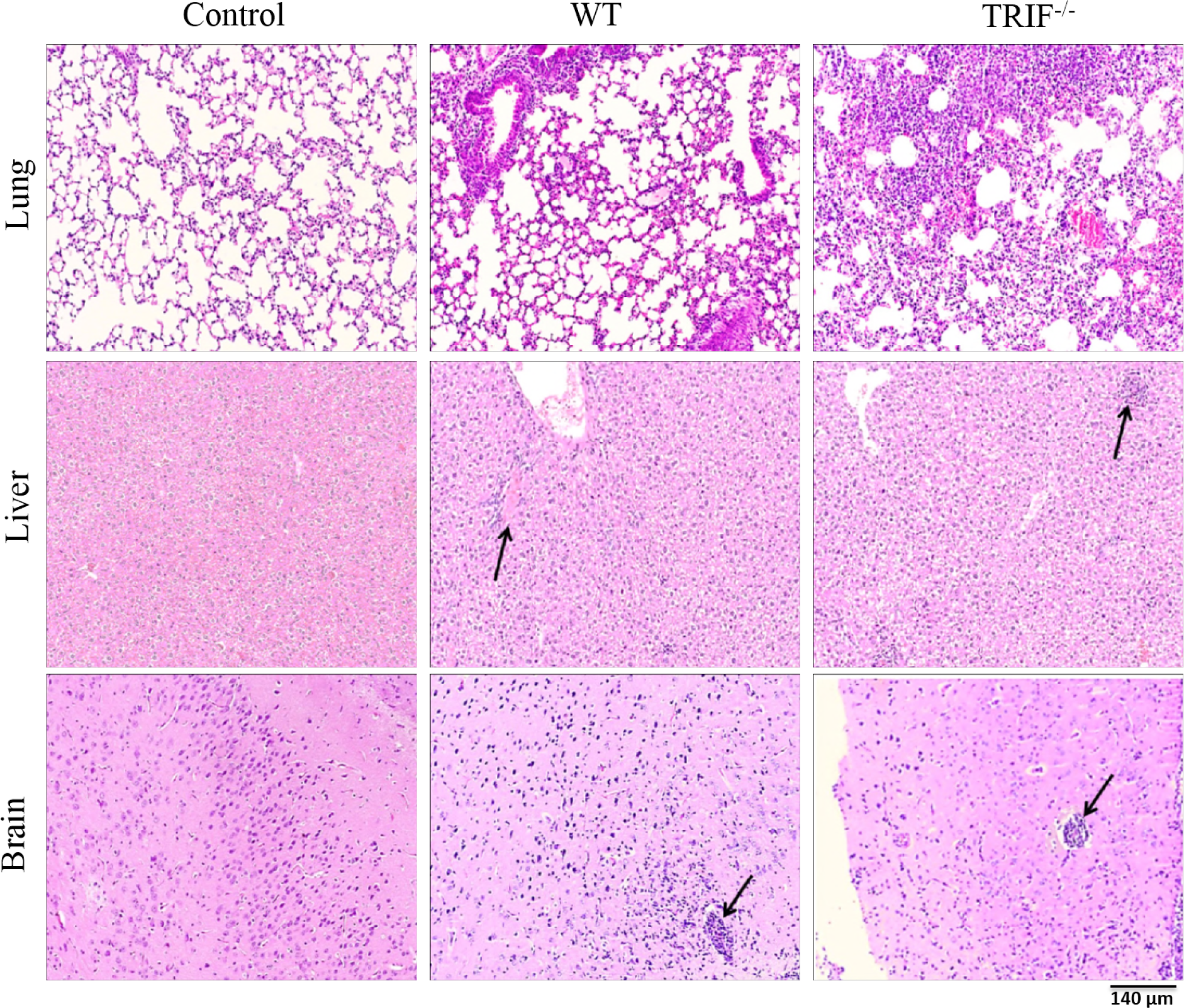
TRIF limits pulmonary inflammation during acute *N. caninum* infection. Representative photomicrographs of histological sections obtained from WT and TRIF^−/−^ (n = 5) mice infected with 5×10^6^ tachyzoites of *N. caninum*. Pulmonary and hepatic tissue samples were collected with 7 days of infection and brain tissue with 30 days of infection. Black arrows indicate areas of inflammatory cellular infiltrate. The slides were stained with hematoxylin-eosin and analyzed under an optical microscope (10x magnification). Values are representative of two independent experiments.

### 2.2 TRIF supports the induction of appropriated Th1 immune profile responses against *N. caninum*

As TRIF^−/−^ mice presented impaired resistance to the infection, shown by decreased survival rates, along with higher parasite burden and tissue injury, we investigated whether the stimulation of adaptive immune responses, based on the production of a proper profile of chemokines, cytokines and nitric oxide, would also be compromised in TRIF^−/−^ mice during the acute phase of infection by *N. caninum*. For this purpose, we mapped important soluble factors involved in the activation, regulation and chemoattraction of monocytes, lymphocytes and macrophages, in lysed spleen cells obtained from infected WT and TRIF^−/−^ mice during acute phase. The results for the array assay show that TRIF^−/−^ mice fail to produce regular amounts of several chemokines, such as IP-10, JE, CXCL9, CCL5, if compared to the WT (Figure 4). In addition, we found that the concentration of major proinflammatory cytokines such as IL-12p40 (Figure 5A), IFN-γ (Figure 5B) and TNF (Figure 5C) were significantly reduced in the peritoneal fluids and lung homogenates of TRIF^−/−^ mice during acute infection. As expected, *in vivo* nitrate/nitrite production was also restricted in the absence of TRIF. These results indicate that this adapter molecule has a relevant role in the development of appropriated effector immune responses against *N. caninum* infection (Figure 5D).

**Figure 4:**
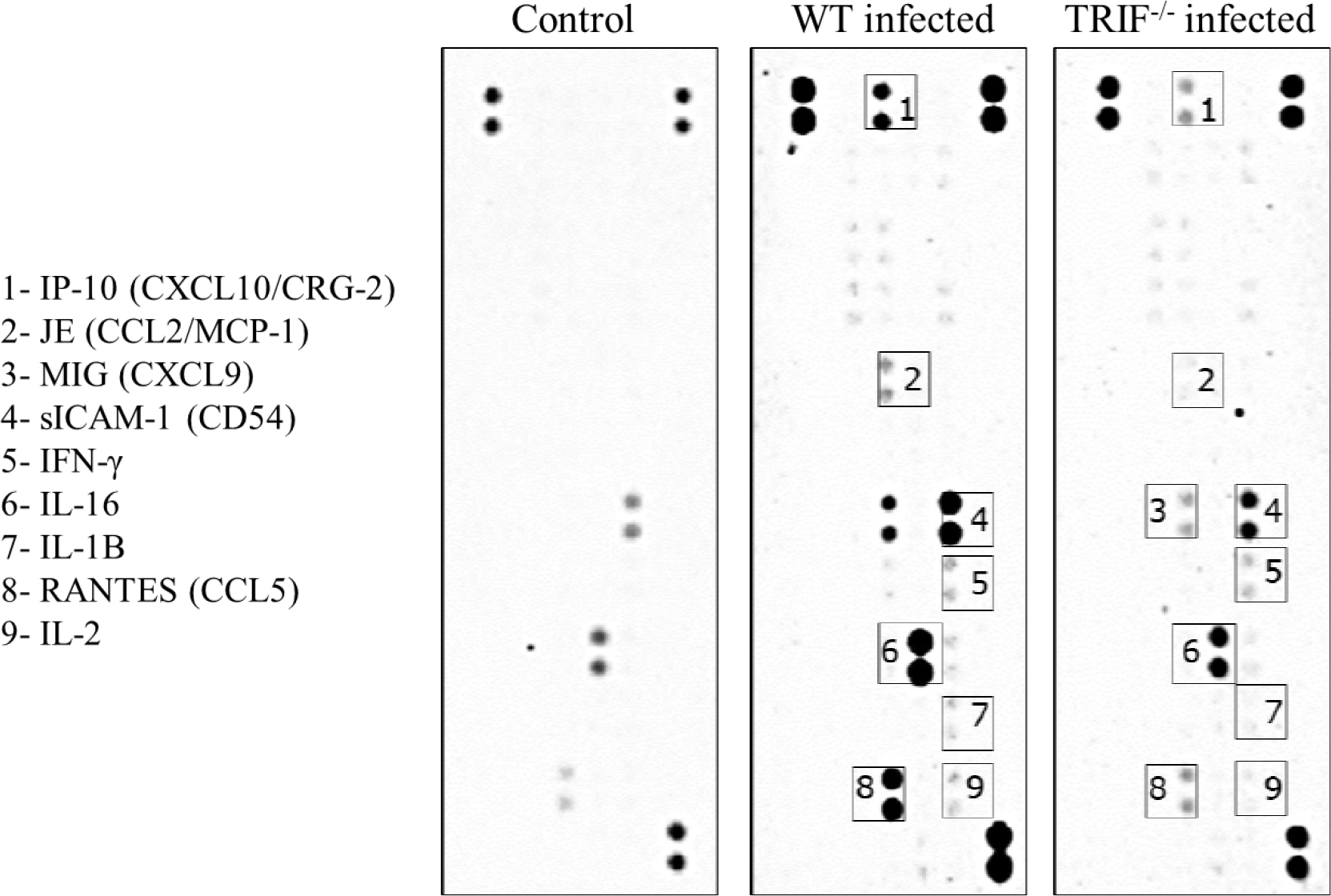
The absence of TRIF impairs adequate production of key cytokines and chemokines during *in vivo* infection. Proteome Profiler Array of lysed spleen cells obtained from WT and TRIF^−/−^ mice 5 days after infection by *N. caninum* (5×10^6^ tachyzoites), showing the expression of different cytokines and chemokines produced by innate and adaptive immune responses.

**Figure 5:**
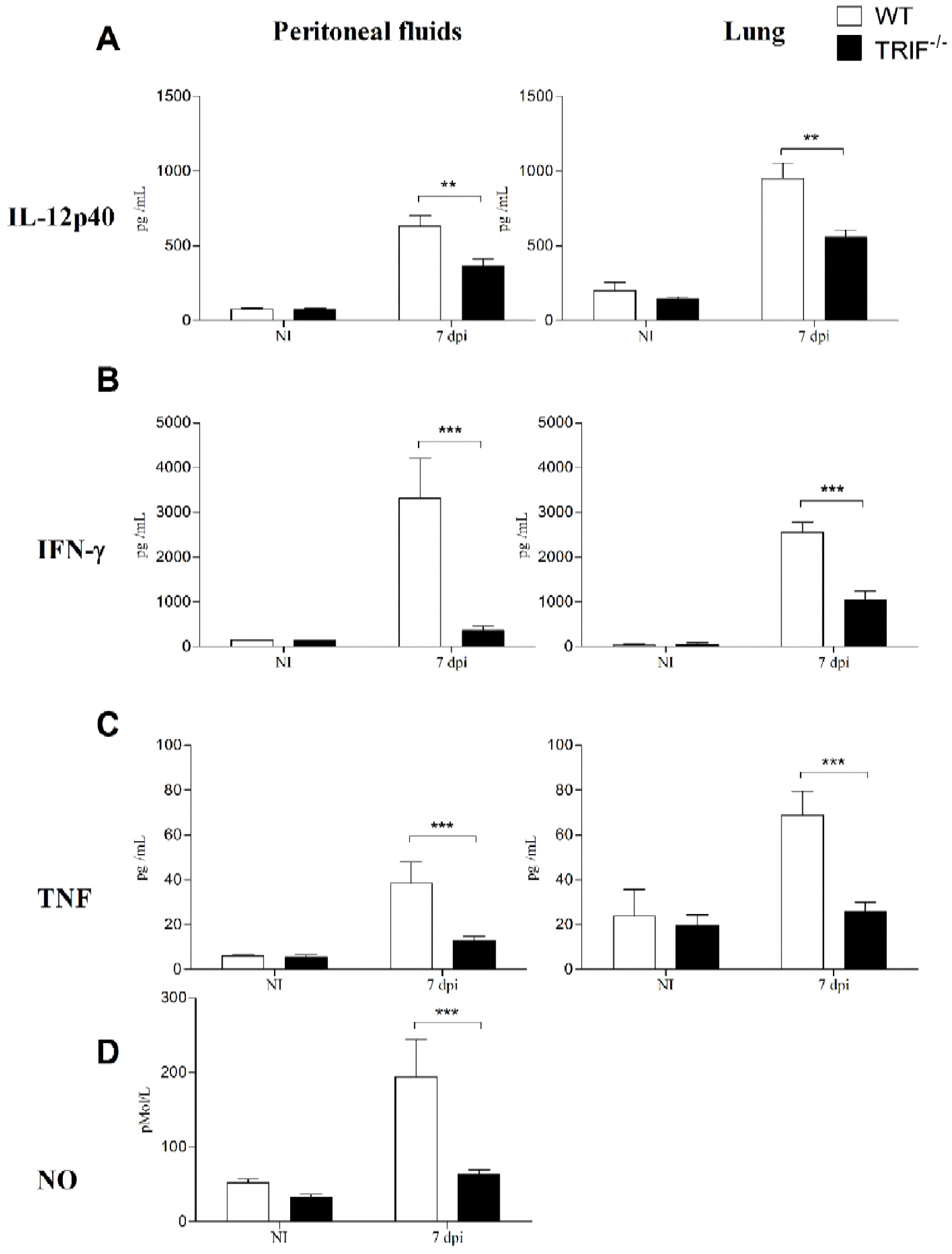
TRIF is partially required for production of pro-inflammatory cytokines and nitric oxide *in vivo*. Quantification of cytokines (**A**- IL-12p40, **B**- IFN-γ and **C**- TNF) production and nitric oxide (**D**) in peritoneal fluids and lungs from WT and TRIF^−/−^ mice (n = 5), during the acute phase of infection by *N. caninum* (7 dpi, 5×10^6^ tachyzoites/mice). The results were expressed as mean ± standard error and were analyzed using the Two-way ANOVA test followed by Bonferroni’s post-test. Values are representative of two independent experiments. (** P <0.01, *** P <0.001).

### 2.3 Differently from *T. gondii*, TLR3 and IRF3 are recruited to the *N. caninum* parasitophorous vacuole (PV)

After analyzing the relevance of the TLR3-TRIF-dependent signaling pathway in the host resistance to *N. caninum*, the next step was to elucidate aspects of the interaction between TLR3 and the parasite, also comparing it to *T. gondii*, which has been previously described as not recognizable to TLR3 (19). First, in order to verify the recruitment of TLR3 in response to *N. caninum* and *T. gondii*, we infected immortalized bone marrow derived macrophages (iBMDMs), transfected with a TLR3-GFP construct, with tachyzoites forms (Figure 6) or total RNA extracted from both parasites (Figure 7). We observed that the synthesis of TLR3-GFP was enhanced in *N. caninum* infected macrophages, which colocalized with the PV after 24 hours of infection. On the other hand, *T. gondii* did not induce notable expression and foci formation of this receptor. In agreement, the exposure of iBMDMs to *N. caninum* total RNA recruited the TLR3-GFP dispersed in cytoplasm to form agglomerates inside the cell, while *T. gondii* RNA was again not able to induce recruitment of this receptor. When a transfecting reagent was used, in order to force RNA uptake by iBMDMs, no formation of TLR3-GFP foci was observed, however it was notable that TLR3 was abundantly expressed in cells exposed to *N. caninum* RNA, while GFP expression in cells stimulated with *T. gondii* RNA remained basically unaltered.

**Figure 6:**
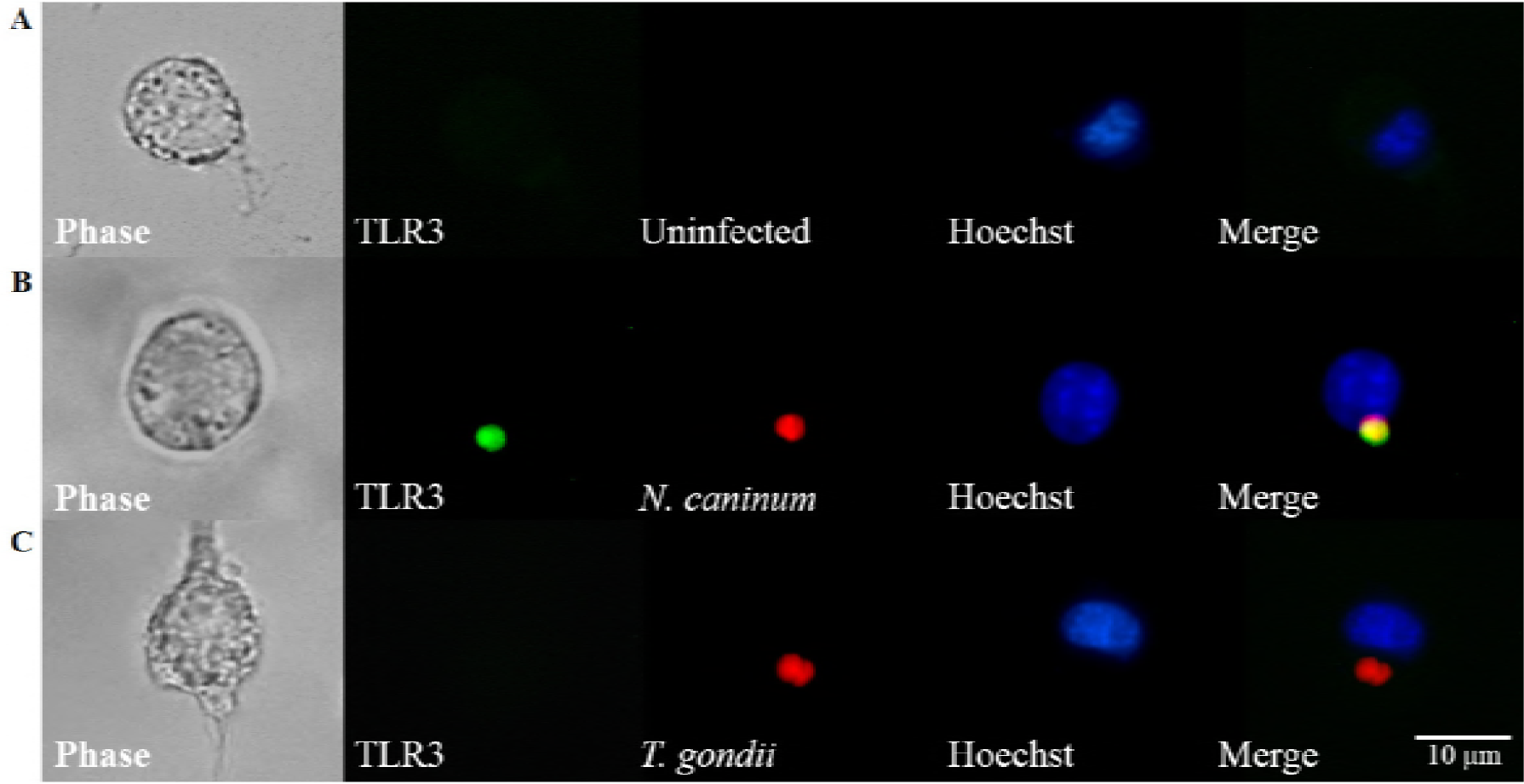
Differential induction of TLR3 recruitment to the PV after infection with *N. caninum* and *T. gondii*. Immortalized murine macrophages transfected with a TLR3-GFP construct after 24 hours of infection with *N. caninum* (Alexa 546) and *T. gondii* (RH-RFP+ strain). (**A**) Uninfected TLR3-GPF+ macrophages. (**B**) TLR3-GPF^+^ macrophages infected with *N. caninum* tachyzoites. (**C**) TLR3-GPF^+^ macrophages infected with *T. gondii* tachyzoites. Images are representative of three independent experiments.

**Figure 7:**
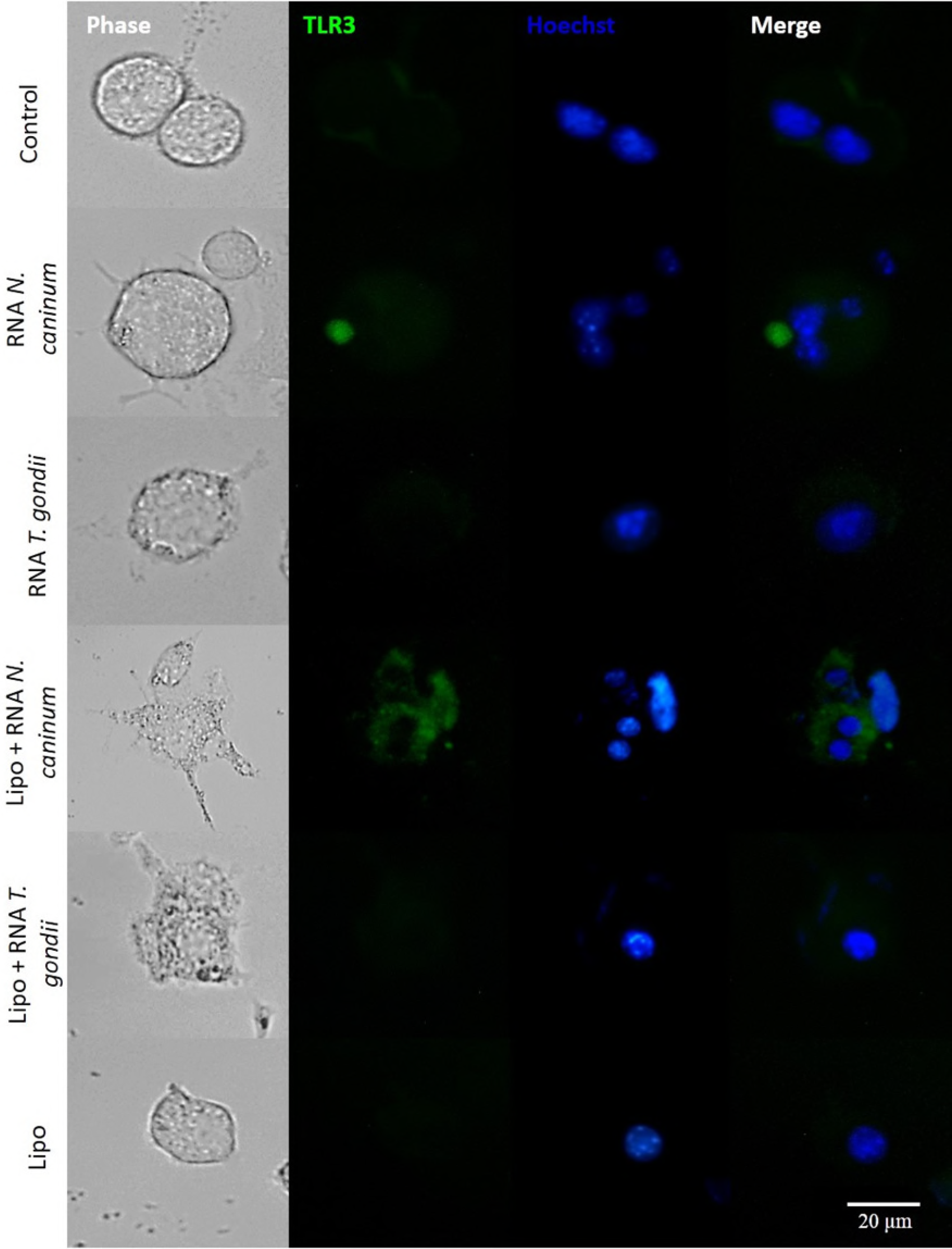
*Neospora caninum* RNA triggers TLR3 expression and agglomeration around the PV. Immortalized murine macrophages transfected with a TLR3-GFP construct were plated and stimulated with 1μg of *N. caninum* or *T. gondii* total RNA, with or without transfecting reagent (lipofectamine - Lipo), for 24 hours. Images are representative of two independent experiments.

In the same manner and still trying to comprehend the mechanisms involved in the recruitment of pathway components to the PV, we transfected macrophages with IRF3-GFP constructs, a transcription factor responsible for the induction of IFN-α and IFN-β. These cells were infected with tachyzoites of *N. caninum* and *T. gondii* for 24 hours and evaluated by fluorescence microscopy (Figure 8). The results showed that infection by *N. caninum* strongly induces increased expression of IRF3 with formation of aggregates around the parasite. Conversely, *T. gondii* was not able to activate IRF3, corroborating with the results previously presented in relation to TLR3 recruitment.

**Figure 8:**
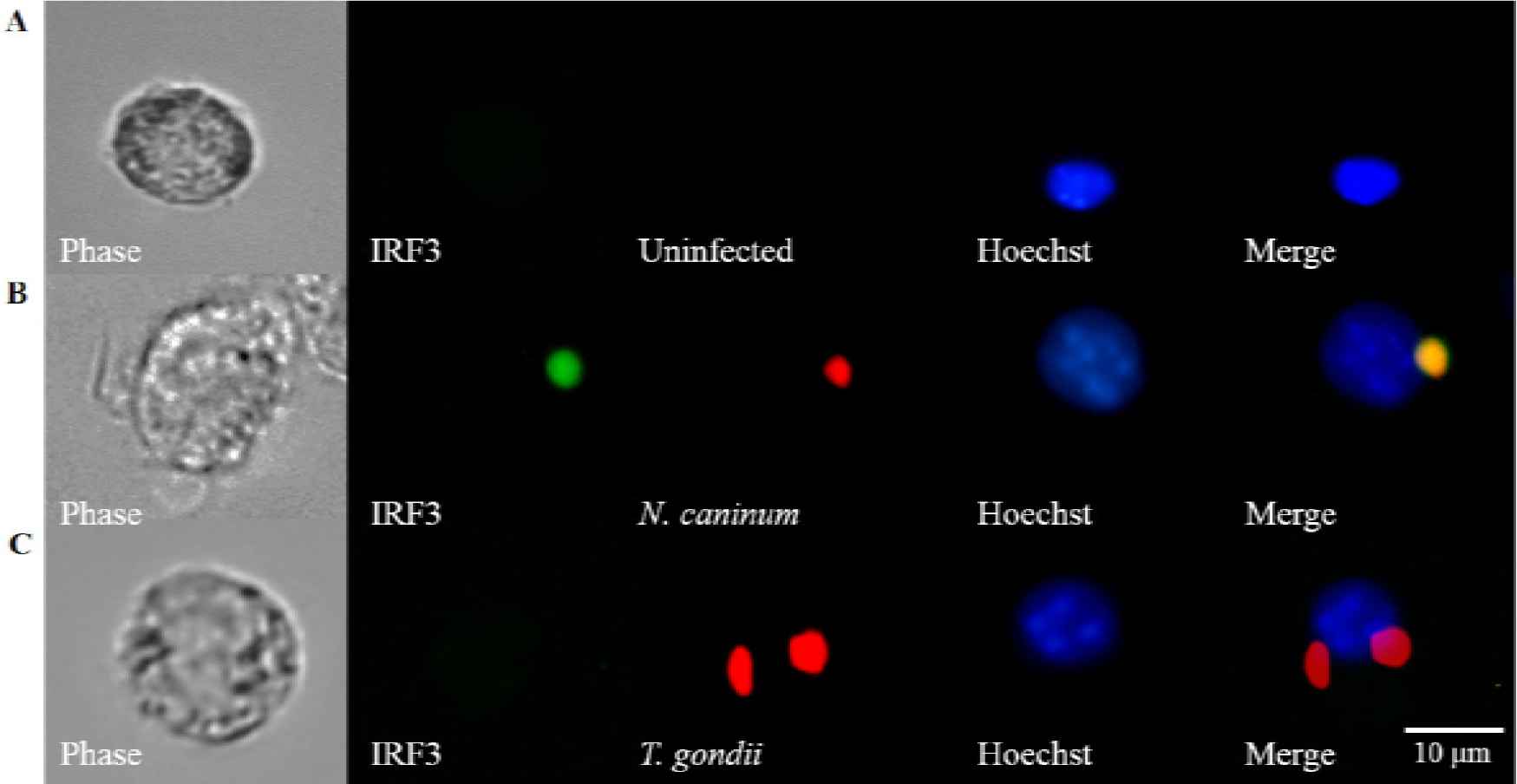
Differential induction of IRF3 recruitment to the PV after infection with *N. caninum* and *T. gondii*. Immortalized murine macrophages transfected with a IRF3-GFP construct after 24 hours of infection with *N. caninum* (Alexa 546) and *T. gondii* (RH-RFP+ strain). (**A**) Uninfected IRF3-GPF^+^ macrophages. (**B**) IRF3-GPF^+^ macrophages infected with *N. caninum* tachyzoites. (**C**) IRF3-GPF^+^ macrophages infected with *T. gondii* tachyzoites. Images are representative of two independent experiments.

### 2.4 TRIF is responsible for the induction of type I IFN in macrophages infected with *N. caninum*, which are shown to be relevant modulators of key IL-12 production

We have demonstrated that protozoan *N. caninum* is able to activate the signaling pathway initiated by TLR3, inducing the recruitment of the receptor and related transcription factor to the PV. We also know from the literature that this signaling pathway is TRIF-dependent and results in type I IFN (IFN-α and IFN-β) production, possibility that we now investigated during infection of fresh BMDMs. Initially, to assess if *N. caninum* was able to modulate the expression of TRIF during the infection, WT BMDMs were exposed to live *N. caninum* tachyzoites, or stimulated with LPS as positive control, and the messenger RNA codifying this adaptor protein was quantified after 6 hours of infection. For this experiment, we observed that TRIF expression increased over 10-fold in *N. caninum* infected BMDMs, if compared to naïve cells (Figure 9A). A similar increase in transcription was observed in the follow up experiments, involving the quantification of IFN-α and IFN-β expression in infected BMDMs derived from WT and TRIF^−/−^ mice. However, in the absence of TRIF, this expression was severely compromised. IFN-α expression was reduced in approximately 40% in infected TRIF^−/−^ BMDMs (Figure 9B), while differences in IFN-β transcript levels were even higher, with TRIF^−/−^ BMDMs inducing over 4-fold reduced expression of this cytokine, if compared to WT BMDMs in the presence of the parasite (Figure 9C).

**Figure 9:**
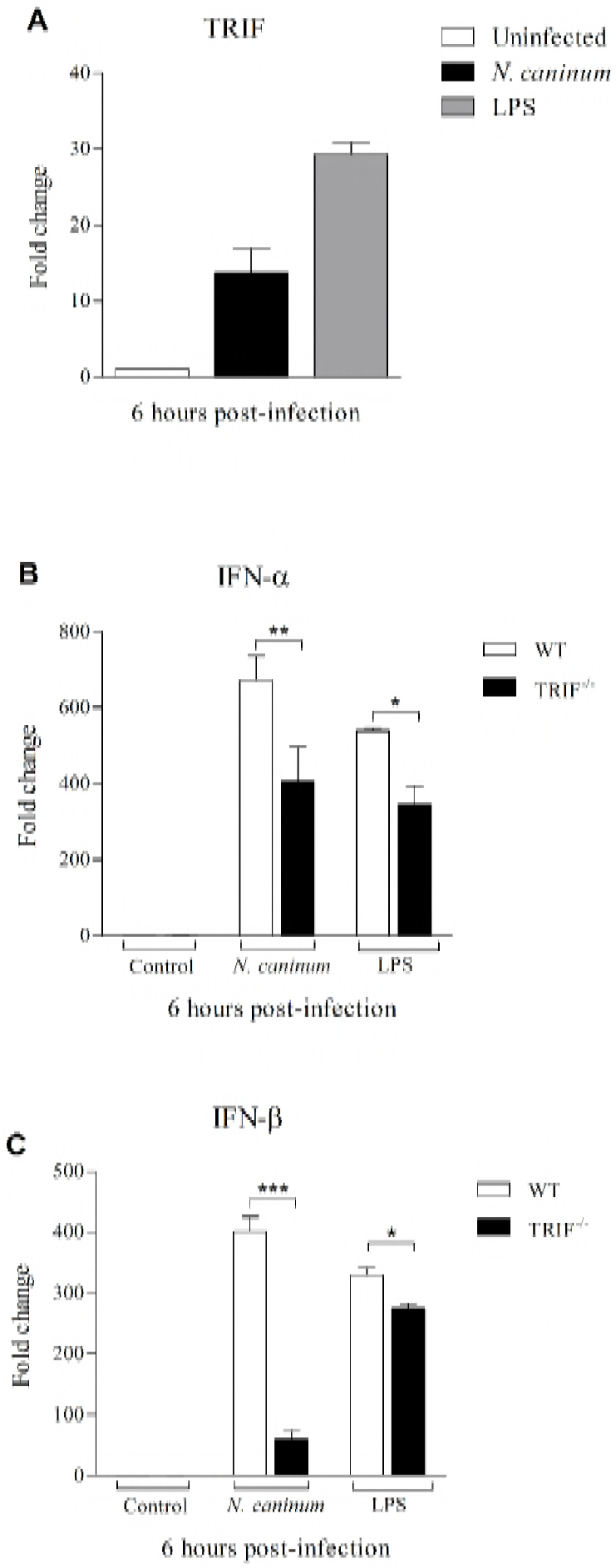
TRIF is upregulated in macrophages after infection and coordinates the gene expression of type I IFNs. A) Quantitative TRIF expression in bone marrow derived macrophages from WT mice infected with *N. caninum* (MOI 0.5) or stimulated with LPS (1 ug/ml). B) IFN-α and C) IFN-β gene expression in bone marrow derived macrophages from WT and TRIF^−/−^ mice infected with *N. caninum* tachyzoites (MOI 0.5) or stimulated with LPS. Data analysis was performed considering the expression results obtained by uninfected macrophages as reference values and are shown as fold change (relative increase of final value in relation to reference value). The results were expressed as mean ± standard error, were analyzed using the Two-way ANOVA test followed by Bonferroni’s post-test, and were representative of three independent experiments. (*P <0.1 ** P <0.01, *** P <0.001).

Our last step was to evaluate whether the TLR3-TRIF pathway and its known products (type I IFNs mainly) would affect Th1 induction by macrophages. As previously described by Mineo and colleagues (12), MyD88 is a crucial adapter molecule in inducing an effective immune response against the parasite due its pivotal role in IL-12 production, which by its turn is the main IFN-γ inducing element in the immune system by polarized Th1 lymphocytes. With that intent, we performed experiments to elucidate whether the TLR3-TRIF pathway would directly affect this essential mechanism in host protection during *N. caninum* infection. For this, we analyzed the production of IL-12p40 by WT, TLR3^−/−^, TRIF^−/−^, and MyD88^−/−^ infected macrophages. In this set of experiments, we observed that TLR3^−/−^ and TRIF^−/−^ BMDMs failed to produce regular concentrations of IL-12p40, if compared to WT cells. As expected, in the absence of MyD88, no production of the analyzed cytokine was detected (Figure 10A).

**Figure 10:**
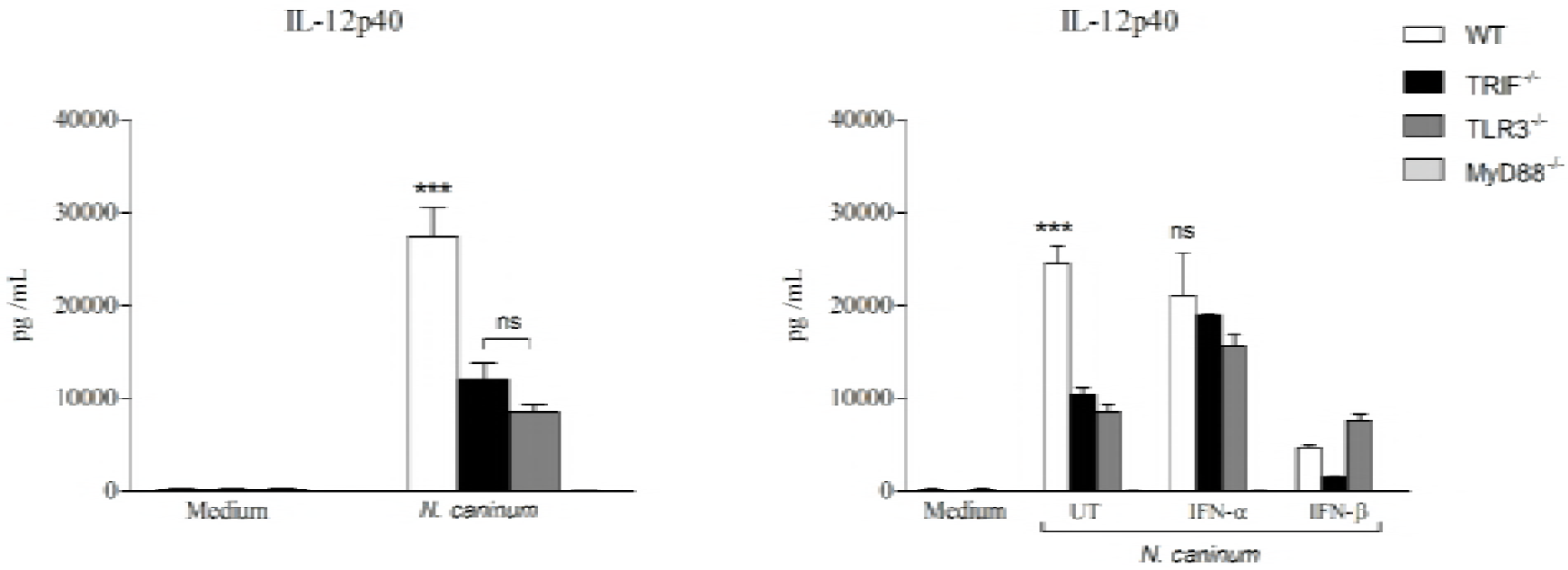
The production of IL-12 is enhanced by TLR3-TRIF-IFN-α pathway during *N. caninum* infection. **(A)** Quantification of IL-12p40 ELISA in the culture supernatants of BMDMs obtained from WT, TLR3^−/−^, TRIF^−/−^ and MyD88^−/−^ mice infected with *N. caninum* tachyzoites (MOI 0.5). **(B)** Quantification of IL-12p40 production in the culture supernatant of BMDMs derived from WT, TLR3^−/−^, TRIF^−/−^ and MyD88^−/−^ mice treated or not with recombinant cytokines IFN-α and IFN-β (10 U/ml) and infected with *N. caninum* tachyzoites (MOI 0.5). Cytokine concentrations were determined after 24 hours of infection. Values are expressed as mean ± standard error. (*** P <0.001, ns: not statistically significant). The data were analyzed using the Two-way ANOVA test followed by Bonferroni’s post-test and the results are representative of three independent experiments.

In order to establish the mechanism by which TLR3-TRIF contribute to IL-12 production in macrophages, TRIF^−/−^ and WT BMDMs were treated with recombinant IFN-α and IFN-β and infected with *N. caninum* tachyzoites for subsequent determination of IL-12 levels. According to the results, treatment with IFN-α was able to completely reestablish IL-12p40 production in TLR3^−/−^ and TRIF^−/−^ BMDMs, while IFN-ß had an overall inhibitory effect on the production of IL-12p40 in all macrophage lineages tested (Figure 10B). None of the treatment with recombinant cytokines were able to induce IL-12p40 in MyD88^−/−^ BMDMs.

## 3. Discussion

*Neospora caninum* is the obligatory intracellular parasite that causes neosporosis and has medical-veterinary importance mainly by causing neuromuscular paralysis in dogs and abortion in cattle, with significant economic impact on countries that produce meat and dairy products. In order to prevent the spread of the infection and its clinical consequences, the understanding of the innate immune pathways triggered against this parasite, as well as the participation of parasite molecules that actively interfere in host resistance to the infection, are crucial (3).

In this work, we investigated how TLR3-TRIF innate recognition pathway assists the host response against the infection by *N. caninum*. Initially, we aimed to evaluate the importance of the this pathway in survival of *N. caninum* infected hosts, using MyD88^−/−^ mice as a comparison, since this molecule is central for survival of the host during neosporosis (12). In our study, TLR3^−/−^ and TRIF^−/−^ mice presented decreased survival rates after infection, although part of the experimental groups survived – differently from MyD88^−/−^ mice groups that fully succumbed to the *N. caninum* challenge. These experiments demonstrate that the TLR3-TRIF pathway has an indirect participation in host resistance to infection, in a secondary role to MyD88 in resistance to the infection.

As observed during *in vivo* experiments, our results showed that TRIF is also related to the regulation of effector host immune responses, since TRIF^−/−^ mice presented higher parasite burden associated with low concentrations of IFN-γ and NO, which are essential for parasite elimination. Nonetheless, intense tissue inflammation was observed in the lungs of TRIF-deficient mice with significant loss of pulmonary alveoli structure, characterizing the severity of the infection in the absence of key immune factors. Hsia and collaborators demonstrated that TRIF^−/−^ mice presented higher lung injuries due to impaired ability of lung dendritic cells to support the polarization of the immune responses to a Th1 profile (20). Previous work indicate that iNOS (nitric oxide synthase 2) is absent and NO synthesis is totally impaired in TRIF deficient murine peritoneal cells stimulated with poly: IC or LPS (21). There is also evidence that TRIF participates not only with NO production, but also with the induction of reactive oxygen species (ROS) in macrophages via TLR3 receptor (22). In addition, we have observed thoroughly that the absence of this adapter molecule also leads to an impaired production of IL-12p40, TNF and important chemokines, indicating that TRIF pathway acts in the immune control of the acute phase of disease through the potentiation of specific Th1 responses, coordinated by MyD88.

From these results, we decided to explore the mechanism by which *N. caninum* interacts with the TLR3-TRIF pathway. Despite its phylogenetic similarity with *T. gondii, N. caninum* has many distinct biological characteristics, especially those regarding it’s interaction with the host cell. Our results demonstrate that one of these differences consists in the ability of *N. caninum* to induce the innate recognition pathway dependent on the TLR3 receptor, which is well characterized for recognizing viral nucleic acids. We have shown that the RNA of the protozoan *N. caninum* triggers the induction of TLR3, recruiting this receptor dispersed in cytoplasm for the formation of an active agglomerate around the PV. Similar results were found by Beiting and colleagues, demonstrating that protozoan parasites are capable of inducing type I IFN responses and that *N. caninum* performs this activation in a higher magnitude than *T. gondii* (23). We also evaluated the behavior of the transcription factor IRF3 by *N. caninum*. Again, a similar behavior was observed, with the concentration of IRF3 around the PV. Beiting and collaborators did demonstrate that *N. caninum* is also able to induce increased expression of IRF7, another transcription factor involved in antiviral responses (23).

Regarding the ability of the TLR3-TRIF pathway to induce IFN-α and IFN-β, we have here confirmed this mechanism in the *Neospora* model. Many examples of this induction may be found in the literature: in a study that used mesothelial cells, which act as a protective barrier against invasive pathogens, it was observed that stimulation with poly I:C, a TLR3 agonist, induces the expression of IFN-β mRNA in WT cells, however, this induction was significantly reduced in TRIF deficient cells (21); In other infection models, such as the bacteria *Listeria monocytogenes* and *Chlamydia muridarum*, it has been observed that macrophages deficient in TRIF also present impaired IFN-β expression, demonstrating again the strict relation of this adapter molecule to this cytokine (24, 25). On the other hand, it is well known that the development of a suitable Th1 immune response is considered of primary importance for the protection against intracellular parasites such as *N. caninum*. The new information to us was the link between type I IFN production and the establishment of a robust and classical Th1 profile, mediated through the partly TLR3-TRIF-dependent production of IL-12. This Th1 induction pathway is not usually assessed by the scientific community, but it has been previously shown that LPS stimulation in the absence of TRIF significantly compromises the production of proinflammatory cytokines as IL-12p40 and TNF (26). Further, we also show the direct effect of type I IFN within this context, as IFN-∝ completely rescued IL-12 production in TRIF^−/−^ macrophages. Similarly, it was demonstrated that IFN-α treatment induced high levels of IL-12 secreted by these cells (27). In addition, it is known that the action of IFN-α and IL-12 together can recruit macrophages and NK cells and elevate the levels of other proinflammatory cytokines, generating a positive feedback loop (28, 29).

Together, our results shown that, unlike *T. gondii, N. caninum* is able to induce the activation of the TLR3-TRIF-IRF3-IFN-α/β signaling pathway, classically known to act on viral infections. In addition, we have demonstrated that TRIF is required for proper resistance against the infection induced by *N. caninum*, enhancing Th1 immune responses against the parasite, which confers increased survival, reduced parasite burden and decreased inflammatory lesions to the host. Based on these findings, future work should be directed to prospecting the actual TLR3 agonist contained within *N. caninum*, which may be used for new prophylactic and/or therapeutic approached aimed against neosporosis or other conditions that require Th1 boosting.

## 4 Material and methods

### 4.1 Ethics Statement

All experiments with mice were approved by the animal research ethics committee at UFU (Comitê de Ética na Utilização de Animais da Universidade Federal de Uberlândia—CEUA/UFU; protocol number 109/16) and were carried out in accordance with the recommendations in the Guiding Principles for Biomedical Research Involving Animals of the International Council for Laboratory Animal Science (ICLAS), countersigned by the Conselho Nacional de Controle de Experimentação Animal (CONCEA; Brazilian National Consul for the Control of Animal Experimentation) in its E-book (http://www.mct.gov.br/upd_blob/0238/238271.pdf). The institution’s animal facility (Centro de Bioterismo e Experimentação Animal—CBEA/UFU) is accredited by the National’s Commissions in animal experimentation (CONCEA, CIAEP: 01.0105.2014) and biosecurity (CTNBio, CQB: 163/02).

### 4.2 Parasites

*N. caninum* tachyzoites (Nc-Liv isolate) and T. gondii (RH strain) were maintained in confluent monolayers of HeLa cells (CCL-2, ATCC, USA) at 37°C with 5% CO_2_, in RPMI 1640 medium supplemented with glutamine (2mM) and antibiotics/antimycotics solution (Thermo Scientific, Wilmington, EUA). Following cellular disruption, the extracellular parasites were obtained by centrifugation of the supernatant at 800 x g for 10 minutes at 4 °C. The pellet was resuspended in RPMI 1640 and the tachyzoites were estimated in a Neubauer chamber for *in vivo* and *in vitro* experiments.

### 4.3 Animals and experimental infections

Wild type (WT) C57BL/6 mice, along with TLR3, TRIF and MyD88 genetically deficient (TLR3^−/−^, TRIF^−/−^, MyD88^−/−^) littermates, 6-8 weeks old, were obtained and maintained at CBEA/UFU, in individually ventilated cages under controlled conditions of temperature (22-25°C) and light (12h/12h), without restriction of water and food, and were submitted to acute (7 days) and chronic (30 days) infection protocols, as well as the analysis of survival after infection with *N. caninum* tachyzoites.

### 4.4 Quantification of the parasite burden

The parasite burden of peritoneal cells and other sampled tissues was determined by qPCR normalized to glyceraldehyde 3-phosphate dehydrogenase (GAPDH), as previously described (30). Briefly, DNA was extracted from 50 mg of cells or tissue using SDS and proteinase K, and its content was estimated by 260/280 ratio (NanoDrop, Thermo Scientific, EUA). The concentration was adjusted to 40 ng/μl for qPCR assays, which used specific primers designed for the Nc-5 sequence of *N. caninum* and GAPDH listed in Table 1. Parasite burden was estimated through the extrapolation of the number of Nc5 copies in the samples compared to predetermined standard curves with DNA obtained from known counts of *N. caninum* tachyzoites, using dedicated equipment and its software (Step One Plus, Thermo Scientific, EUA).

**Table 1:**
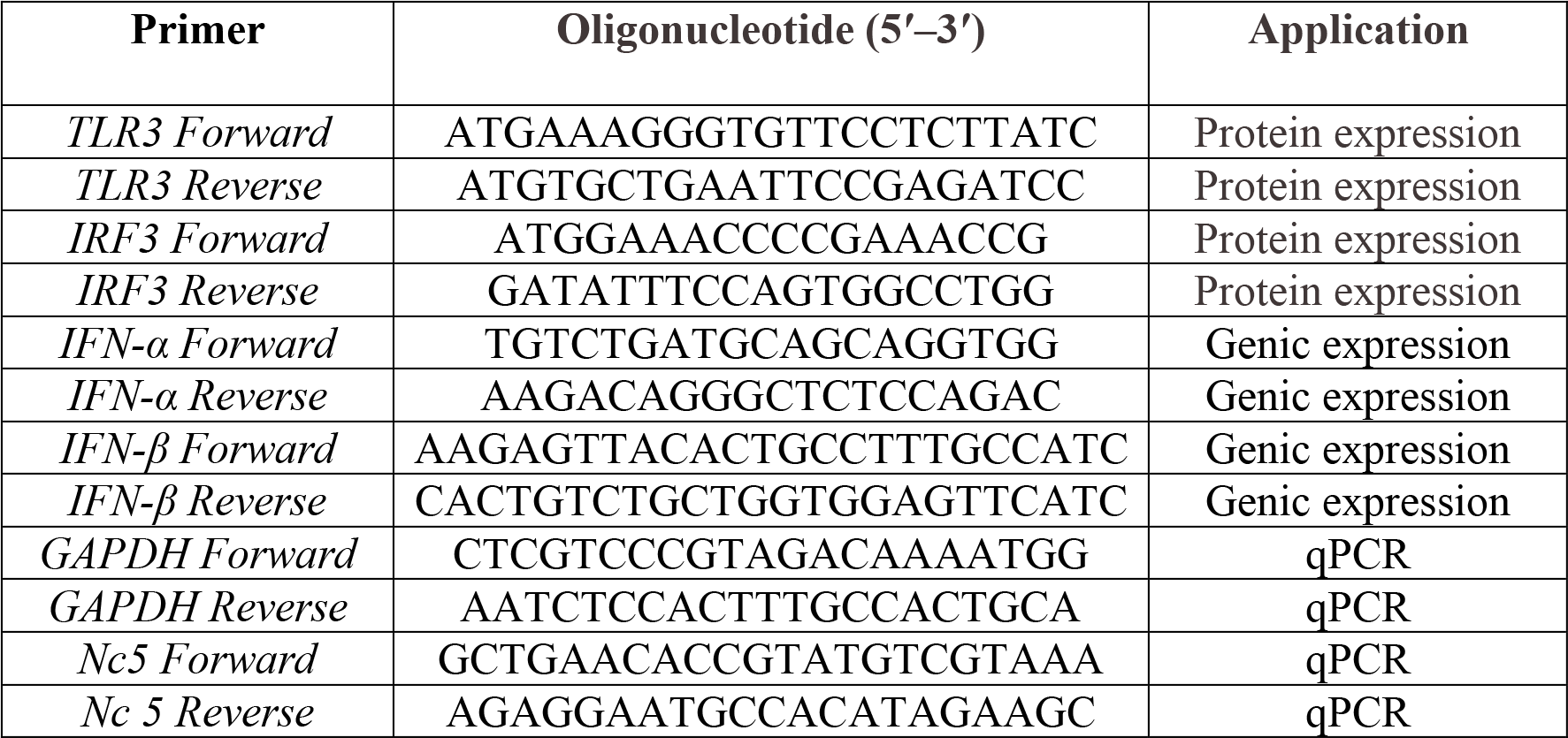
Oligonucleotide primers used in this study

### 4.5 Histological Analysis

Samples of lungs, livers and brains were collected from WT and TRIF^−/−^ mice at different stages of infection and analyzed for inflammatory infiltrates and tissue damage. For this, the samples were fixed in buffered formalin and maintained in alcohol 70% before being submitted to paraffin embedding, ultrathin sectioning and subsequent staining with hematoxylin and eosin (31). The analysis was performed using automated inverted microscope (FSX100, Olympus, Japan).

### 4.6 Cytokine measurement

For the measurement of the cytokine profile and chemokine production, spleens from WT and TRIF^−/−^ mice infected by *N. caninum* (5×10^6^ tachyzoites), were collected after 5 days of infection in RPMI medium. Cell suspensions were washed in medium and treated with lysis buffer (0.16 M NH_4_Cl and 0.17M Tris-HCl, pH 7.5) and resuspended in complete RPMI medium containing 10% FBS. Viable cells were adjusted to 1×10^7^ for the cytokine array by western blotting (Proteome Profiler Array, R&D Systems, Minneapolis, USA) according to the manufacturer’s instructions. In addition, the production of the main pro-inflammatory cytokines (IL-12p40, IFN-γ and TNF) was also measured individually by ELISA in peritoneal fluids and lung homogenates from WT and TRIF^−/−^ mice, infected or not with 5×10^6^ *N. caninum* tachyzoites for 7 days.

For *in vitro* experiments, WT, TLR3^−/−^, TRIF^−/−^ and MyD88^−/−^ macrophages were plated in 96 well plates (2×10^5^cells/well) and incubated for 18 hours at 37°C and 5% CO_2_. After incubation, the BMDMs were treated or not with recombinant cytokines IFN-α and IFN-β (R&D Systems) at 10 U/ml concentration for 6 hours, using LPS (1μg/ml) as positive control. The cells were infected with viable *N. caninum* MOI 0.5 (Infection Multiplicity = 1 parasite / 2 cells) tachyzoites and incubated for 24 hours. Cell culture supernatant was collected for further quantification of cytokines by ELISA.

The measurement was performed using commercial ELISA kits, conducted according to the protocols recommended by the manufacturer (R&D Systems; BD Biosciences, San Diego, EUA). The final concentration of the cytokines was determined according standard curves with known concentrations of recombinant proteins and the results were expressed as pg/ml, observing the respective detection limits for each assay: IL-12p40 (15.6 pg/ml), IFN-γ (4.1 pg/ml) and TNF (3.7 pg/ml).

### 4.7 Quantification of Nitric Oxide

Quantification of Nitric Oxide (NO) was carried out in the peritoneal fluids of WT and TRIF^−/−^ mice infected with 5×10^6^ *N. caninum* tachyzoites. The assay determines the nitric oxide concentration based on the enzymatic conversion of nitrate to nitrite, followed by its colorimetric detection at 540 nm. Nitrate/nitrite concentration for each sample was estimated by a standard curve following the manufacturer’s instructions (R&D Systems). Detection limit: 0.78 umol/L.

### 4.8 Differentiation of bone marrow derived macrophages (BMDMs), production of immortalized macrophages lineage and Immunofluorescence assay

Bone marrow derived macrophages (BMDMs) were generated from WT, TLR3^−/−^, TRIF^−/−^ and MyD88^−/−^ mice, using conditioned medium, as previously described (32) and the immortalization of BMDMs was performed by the method of infection with J2 virus (33).

For cloning of murine TLR3 and IRF3, total RNA was extracted by TRIzol from brain of wild type mice and the cDNA was synthesized using the GoScript ™ Reverse Transcription System (Promega, Madison, USA), according to the manufacturer’s instructions. The PCR gene amplification was performed using specific primers for TLR3 and IRF3 listed in Table 1. The resulting PCR products were cloned in mammalian expression vector pcDNA 3.1(+) encoding GFP (Invitrogen-Life Technologies, USA) and the plasmid was linearized with ScaI-HF for stable transfection in 10^6^ immortalized murine macrophages plated in 24-well plates, using LipofectamineTM 2000 (Invitrogen, Carlsbad, CA, USA). The transfection efficiency was monitored under fluorescence microscopy after 72 hours, and the positive cells were cloned and drug selected (Geneticin, G418; Hyclone Laboratories). After cloning, cells were plated at 1×10^6^ /well in 24-well plates and then infected with *N. caninum* or *T. gondii* tachyzoites (MOI 1) for 24 hours. For *N. caninum* labelling we performed an immunofluorescence reaction as previously described (34, 35) using as secondary antibody Alexa Fluor 546-conjugated anti-mouse (Life Technologies). For infection with *T. gondii* we used a RH_RFP fluorescent strain.

### 4.9 Quantification of gene expression in macrophages

After differentiation, WT and TRIF^−/−^ BMDMs were plated in 24 well plates (10^6^ cells/well) and left at 37°C and 5% CO_2_ for 18 hours. After incubation, the cells were infected with *N. caninum* tachyzoites (MOI 0.5) and left for 6 hours before RNA extraction, cDNA synthesis and subsequent analysis of the expression of TRIF, IFN-α and IFN-β expression by qPCR. RNA extraction was performed with Trizol (Thermo Scientific, EUA) protocol. After this procedure, the purity and yield were determined in spectrophotometer for subsequent cDNA synthesis using GoScript™ Reverse Transcription System (Promega, Madison, EUA). The gene expression assay (qPCR) was performed using SYBR green (Promega) with specific primers listed in Table 1. The gathered data were analyzed by relative gene expression, as previously described (36) using GAPDH as the housekeeping gene. The data was displayed as the fold increase in gene expression, using uninfected WT macrophages as the baseline parameter.

### 4.10 Statistical analysis

Statistical analyzes were performed using a dedicated software (Prism 6.0 GraphPad Software Inc.). The results were expressed as mean ± standard error and the differences were considered statistically significant when p <0.05, as determined by two-way ANOVA with multiple comparison Bonferroni post-tests, T test or Mann Whitney test, depending of the peculiarity of each experimental protocol.

## Acknowledgments

The authors thank Ana Claudia Arantes Marquez Pajuaba, Marley Dantas Barbosa, Murilo Vieira da Silva and Zilda Mendonça da Silva Rodrigues for their technical assistance. This work was supported by Brazilian Funding Agencies (CNPq, FAPEMIG and CAPES).

